# The relationship between reinforcement and explicit strategies during visuomotor adaptation

**DOI:** 10.1101/206284

**Authors:** Olivier Codol, Peter J Holland, Joseph M Galea

## Abstract

The motor system’s ability to adapt to changes in the environment is essential for maintaining accurate movements. During such adaptation several distinct systems are recruited: cerebellar sensory-prediction error learning, success-based reinforcement, and explicit strategy-use. Although much work has focused on the relationship between cerebellar learning and strategy-use, there is little research regarding how reinforcement and strategy-use interact. To address this, participants first learnt a 20° visuomotor displacement. After reaching asymptotic performance, binary, hit-or-miss feedback (BF) was introduced either with or without visual feedback, the latter promoting reinforcement. Subsequently, retention was assessed using no-feedback trials, with half of the participants in each group being instructed to stop using any strategy. Although BF led to an increase in retention of the visuomotor displacement, instructing participants to remove their strategy nullified this effect, suggesting strategy-use is critical to BF-based reinforcement. In a second experiment, we prevented the expression or development of a strategy during BF performance, by either constraining participants to a short preparation time (expression) or by introducing the displacement gradually (development). As both strongly impaired BF performance, it suggests reinforcement requires both the development and expression of a strategy. These results emphasise a pivotal role of strategy-use during reinforcement-based motor learning.

## Introduction

In a constantly changing environment, our ability to adjust motor commands in response to novel perturbations is a critical feature for maintaining accurate performance ^1^. These adaptive processes have often been studied in the laboratory through the introduction of a visual displacement during reaching movements ^2^. The observed visuomotor adaptation, characterized by a reduction in performance errors, was believed to be primarily driven by a cerebellar-dependent process that gradually reduces the mismatch between the predicted and actual sensory outcome (sensory prediction error) of the reaching movement ^1,3,4^ Cerebellar adaptation is a stereotypical, slow and implicit process and therefore does not require the individual to be aware of the perturbation to take place ^5,6^. However, a single-process framework cannot account for the great variety of results observed during visuomotor adaptation tasks ^7^. Specifically, it has recently been shown that several other non-cerebellar learning mechanisms also play a pivotal role in shaping behaviour during adaptation paradigms such as explicit strategy-use ^8,9^ and reward-based reinforcement ^10–12^.

Strategy-use usually consists of employing simple heuristics such as aiming off target in the direction opposite to a visual displacement, to quickly and accurately account for it ^5^. However, this requires explicit knowledge of the perturbation, which in turn usually requires experiencing large and unexpected errors ^8,13–15^ Strategy-use contrasts with cerebellar adaptation in that it is idiosyncratic ^9^, explicit, and can lead to fast adaptation rates ^16^. Critically, cerebellar adaptation takes place regardless of the presence or absence of explicit strategies, even at the cost of accurate performance ^5^.

More recently, another putative mechanism contributing to motor adaptation has been proposed, through which the memory of actions that led to successful outcomes (hitting the target) are strengthened, and therefore more likely to be re-expressed. Such reinforcement is considered to be an implicit process, but distinct from cerebellar adaptation in that it doesn’t employ sensory information but task success or failure ^10,11^. To examine this phenomenon, several studies employed a binary, hit-or-miss feedback (BF), paradigm which promotes reinforcement over cerebellar processes ^11,12,17^. For example, in one study, participants receiving only binary feedback following successful adaptation expressed stronger retention than participants who had received a combination of visual and binary feedback ^12^. The authors argued this could be due to greater involvement of reinforcement-based process that is less susceptible to forgetting ^12^.

With the multiple processes framework of motor adaptation, the question of interaction between the distinct systems becomes central to understanding the problem as a whole, and it remains an under-investigated question for reward-based reinforcement. In decision-making literature, it has long been suggested that two distinct “model-based” and “model-free” systems interact ^18,19^ and even require communication to be optimal ^20,21^. Interestingly, model-based processes share many characteristics with strategy-use during motor adaptation, in that they are both more explicit, rely on an internal model of the world (strategy-use ^22,23^; model-based decision-making ^24^), and are closely related to working memory capacity (strategy-use ^25,26^; model-based decision-making ^27,28^) and pre-frontal cortex processes (strategy-use^25^; model-based decision-making ^21,29^). On the other hand, the concept of reinforcement in motor adaptation comes directly from the model-free systems described in decision-making literature ^23^, and is often labelled as such. It is more implicit, relies on immediate action-reward contingencies and is thought to recruit the basal ganglia in both cases (visuomotor adaptation ^17^; decision-making ^18^). Despite these interesting similarities, unlike model-based and model-free decision-making, the relationship between strategy-use and reinforcement during visuomotor adaptation paradigms is currently unknown. Evidence of this relationship exists from a recent study which showed participants needed to experience a large reaching error in order to express a reinforcement-based memory ^15^. As suggested before, strategy-use is an explicit process that requires experiencing large errors ^13,14,22^. Thus, is it possible that the formation of a reinforcement-based memory requires, or at least benefits, from some form of strategy-use.

To address this, we first examined the contribution of strategy-use to the reinforcement-based improvements in retention following binary feedback ^12,17^. Secondly, we used a forced reaction time (forced RT) paradigm ^30^ to investigate the importance of being able to express a strategy when encountering binary (reinforcement-based) feedback.

## Results

### Experiment 1: strategic re-aiming occurs during reinforcement-based retention

We first sought to investigate the role of strategy-use in the retention of a reinforced visual displacement memory. In experiment 1, participants made fast ‘shooting’ movements towards a single target (figure 1a). After a baseline block involving veridical vision (60 trials) and an adaptation block (75 trials) where a 20° counter-clockwise (CCW) visuomotor displacement was learnt with online visual feedback (VF), participants experienced the same displacement for 2 blocks (asymptote blocks; 100 trials each) with either only binary feedback (BF group, figure 1b, top) to promote reinforcement, or BF and VF together (VF group, figure 1b, bottom). Following this, retention was assessed through 2 no-feedback blocks (100 trials each), during which both BF and VF were removed. Before these no-feedback blocks, half of the participants were told to “carry on” as they were (“Maintain” group) and the remaining ones were informed of the nature of the perturbation, and to stop using any strategy to account for it (“Remove” group). Thus, there were four groups: BF-Maintain, BF-Remove, VF-Maintain and VF-Remove (N=20 for each group).

**Figure 1:**
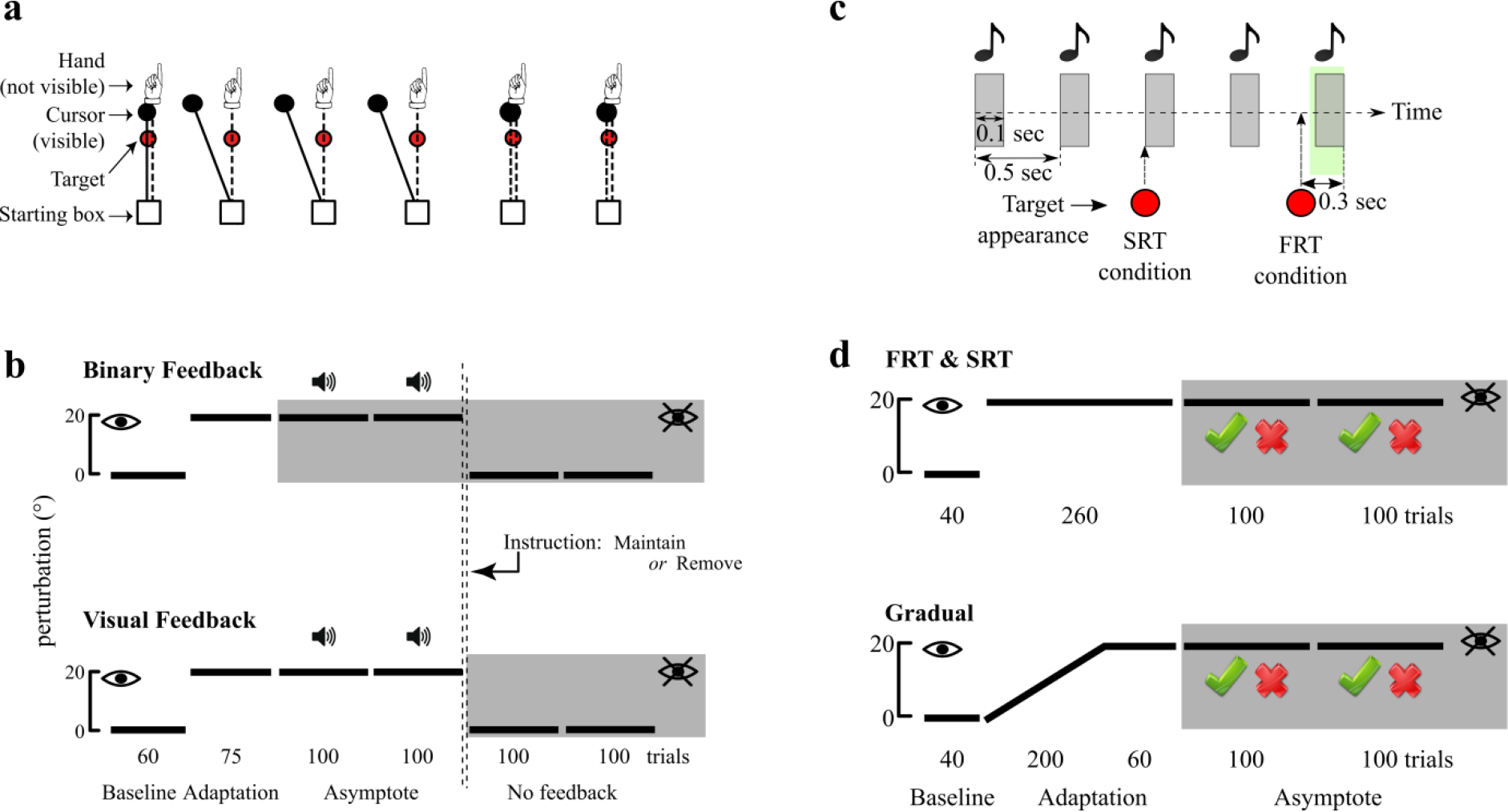
Experimental design. **(a)** Experiment 1: feedback-instruction. Screen display and hand-cursor coupling across each block of the task. **(b)** Feedback-instruction task perturbation and feedback schedule for the BF groups (top) and VF groups (bottom). The white and grey areas represent blocks where VF was available or not available, respectively, as indicated with a crossed or non-crossed eye. Blocks in which hits (with 5° tolerance on each side of the target) were followed by a pleasant sound are indicated with a small speaker symbol. The y-axis represents the value of the discrepancy between hand movement and task feedback. The double dashed vertical lines represents the time point at which “Maintain” or “Remove” instructions were given. The number of trials and names for each block are indicated at the bottom of each schedule. **(c)** Experiment 2: forced RT. Schedule of tone playback and target appearance before each trial during the forced RT task (SRT and FRT conditions). Participants were trained to initiate their reaching movements on the last of a series of five 100 ms-long tones played at 0.5 sec intervals. The green area represents the allowed movement initiation timeframe, and the red dots represent target onset times for each condition. The grey areas represent the tones. **(d)** Forced RT task perturbation and feedback schedule for the SRT and FRT groups (top) and for the Gradual group (bottom). Grey areas represent blocks without VF. The green tick and the red cross represent binary feedback cues for a hit (5° tolerance on each side of the target) and miss, respectively. The white and grey areas represent blocks in which VF was available or not available, respectively, as indicated with a crossed or non-crossed eye, and the y-axis represents the value of the discrepancy between hand movement and task feedback. The number of trials and names for each block are indicated at the bottom of each schedule. BF: binary feedback; VF: visual feedback; RT: reaction time; SRT: slow reaction time; FRT: fast reaction time.

Group performance is shown in figure 2a. All groups showed similar baseline performance (figure 2b; H(3)=4.59 p=0.20; see Methods for detailed information on statistical analysis), and had fully adapted to the visuomotor displacement prior to the asymptote/reinforcement blocks (average reach angle in the last 20 trials of adaptation, figure 2c; H(3)=2.56 p=0.46). Interestingly, at the start of the first asymptote block, participants in both BF groups showed a dip in performance, effectively drifting back toward baseline before adjusting back and returning to plateau performance. This “dip effect” was completely absent in the VF groups. Therefore, success rate was compared independently across groups in the first 30 trials (figure 2d) and the remaining 170 trials (figure 2e) of the asymptote block. Both BF groups exhibited lower success rates than the VF groups in the early asymptote phase (H(3)=46.79, p<0.001, Tukey’s test p<0.001 for BF-Maintain vs VF-Maintain and vs VF-Remove, and for BF-Remove vs VF-Maintain and vs VF-Remove). This was also seen in the late asymptote phase (H(3)=31.29, p<0.001, Tukey’s test p<0.001 for BF-Maintain vs VF-Maintain and vs VF-Remove, and for BF-Remove vs VF-Maintain and vs VF-Remove), although performance greatly improved for both BF groups compared to the early phase (Z=3.692 and Z=−3.81 for BF-Remove and BF-Maintain, respectively, p<0.001 for both). This dip in performance has previously been observed independently of our study when switching to BF after a displacement is abruptly introduced ^12^. Finally, no across-group difference in RTs or movement duration was found during the asymptote blocks (Supplementary figure S1b, c).

Participants then performed a series of 2 no-feedback blocks. Similar to Shmuelof et al., ^12^ we assessed retention by looking at the last 20 trials of the second block. However, our results are fundamentally the same irrespective of the trials used to represent retention. Overall, the BF-Maintain group showed greater retention relative to all other groups, largely maintaining the reach angle values achieved during the asymptote phase, whereas there was no difference between the other groups (figure 2f; H(3)=27.66, p<0.001, Tukey’s test p=0.001 for BF-Remove vs BF-Maintain and p<0.001 for BF-Maintain vs both VF groups; p=0.6 for BF-Remove vs VF-Remove; p=1 for BF-Remove vs VF-Maintain; p=0.68 for VF-Maintain vs VF-Remove). We therefore replicated previous work which showed that BF led to enhanced retention of a visual displacement when compared to VF ^12^. However, this effect of BF was abolished by asking participants to remove any strategy they had developed (BF-remove). This suggests the increase in retention following BF was mainly a consequence of the greater development and expression of a strategy.

**Figure 2.**
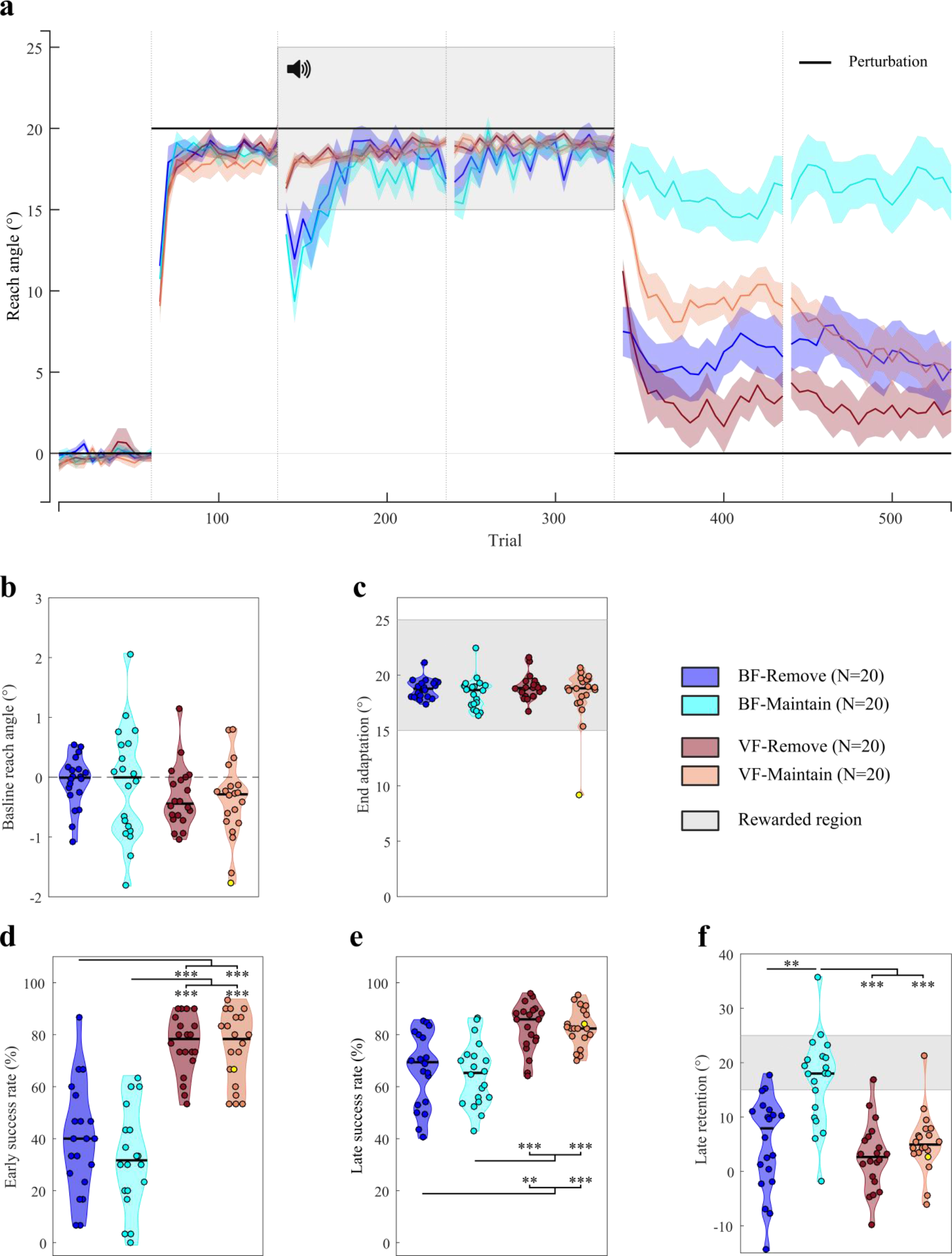
Experiment 1: feedback-instruction. **(a)** Reach angles with respect to target (°) of each group during the visuomotor displacement task. Values are averaged across epochs of 5 trials. Vertical bars represent block limits. The binary feedback consisted of a pleasant sound in the rewarded region. The black solid line represents the hand-to-cursor discrepancy (the perturbation) for all groups across the task. Coloured lines represent group mean and shaded areas represent s.e.m. **(b)** Average reach angle of participants during baseline. **(c)** Average reach angle during the last 20 trials of the adaptation phase. The shaded area represents the region to be rewarded in the subsequent asymptote phase. **(d)** Success rate (%) during the first 30 trials of the asymptote phase. **(e)** Success rate during the remainder of the asymptote phase (i.e. trial 31 to 200 of asymptote blocks). **(f)** Average reach angle during the last 20 trials of the no-feedback (retention) phase. Each dot represents one participant. The yellow dot represents the same participant across all plots, who expressed atypical end adaptation reach angle values; however this was not seen across the other variables. For the distribution plots, horizontal black lines are group medians and the shaded areas indicate distribution of individual values. BF: binary feedback; VF: visual feedback. *** p<0.001, ** p<0.01

### Experiment 2: re-aiming is necessary for maintaining performance under binary feedback

If this conclusion from our first experiment is correct, then successful asymptote performance under BF only should be dependent on the ability to develop and express a strategy. Therefore, in experiment 2 we restricted participant's capacity to use a strategy by using a forced RT adaptation paradigm ^30–32^ (figure 1c). Specifically, two groups adapted to a 20° CCW visuomotor displacement by performing reaching movements to 4 targets (figure 1d), with the amount of available preparation time (i.e. time between target appearance and movement onset) being restricted. A first group was allowed to express slow RTs (SRT; RT constraints were 870 to 1000 ms after target onset; N=10), while the second group was only allowed very fast RTs (FRT; 130 to 300 ms; N=10; figure 1c and Supplementary figure S2a). The latter condition has been shown to prevent time-demanding strategy use such as mental rotations necessary to express re-aiming in reaching tasks ^30,32,33^. Critically, this paradigm only prevented expression of re-aiming, but not strategy development. Therefore, to ensure any between-group difference was task-dependent and not related to inter-individual differences in awareness or understanding of the task, we explained in detail the nature of the perturbation and the optimal strategy to counter it. In addition, a third condition was designed in which participants were kept unaware of the visual displacement by introducing the perturbation gradually^13,15^ (N=10; figure 1d, bottom), and were not informed of any optimal strategy to employ. Participants in this group were given no RT constraint whatsoever. Finally, it should be mentioned that a large portion of participants in the Gradual group reported noticing a slight perturbation by the end of the adaptation block when informally asked after the experiment. However, they underestimated its amplitude significantly at best, reporting effects of the order of 5°. Nevertheless, for the sake of simplicity we will qualify this group as “unaware”, although we hereby acknowledge they reported very partial, reduced awareness of the perturbation.

During baseline, average reach direction was similar for all groups (figure 3b; H(2)=0.45, p=0.79). To examine whether the FRT and SRT groups displayed different rates of learning during adaptation, we applied an exponential model to each participant’s adaptation data. Note, this was not done for the gradual group whose adaptation rate was restricted by the incremental visuomotor displacement. Surprisingly, we found no significant difference between the FRT and SRT group’s learning rates (U=74; p=0.34; Supplementary figure S2b). Indeed, one would expect the SRT group to express faster learning since they can express strategies to account for the perturbation ^16,30,32,34^. This is most likely a consequence of the small size of the perturbation encountered (i.e. 20°), which leaves less margin for strategic re-aiming ^34–36^. At the end of the adaptation block, all groups adapted successfully, with no significant difference in reaching direction (figure 3c; H(2)=2.34, p=0.31). However, despite the lack of statistical significance, the mean reach direction for the FRT group was slightly under 15° (mean: 14.87°), which represents the limit of the reward region in the subsequent block. We discuss the implications of this later.

**Figure 3.**
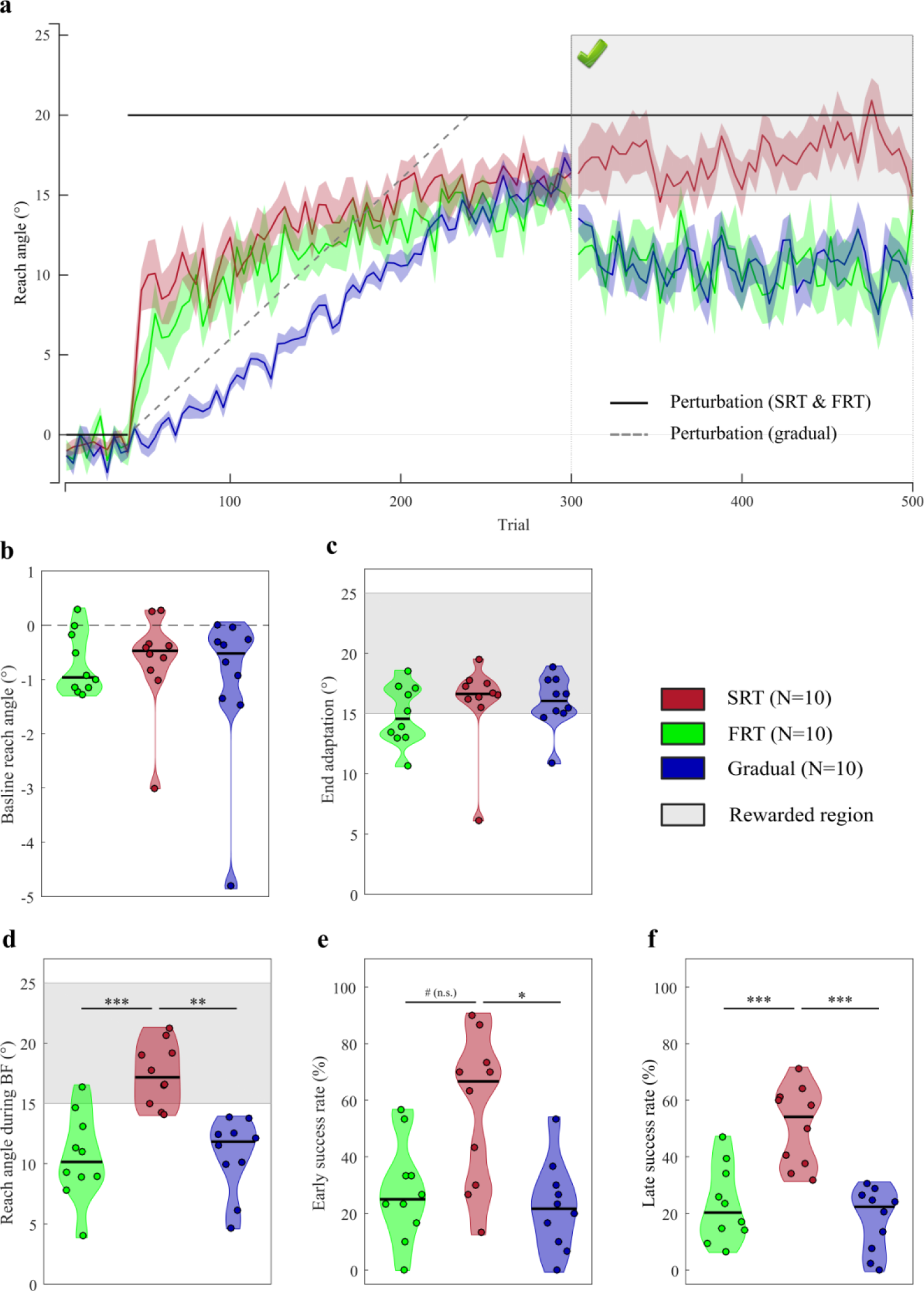
Experiment 2: forced RT. **(a)** Reach angles with respect to target (°) of each group during the visuomotor displacement task. Values are averaged across epochs of 4 trials. Vertical bars represent block limits. The binary feedback consisted of a large green tick displayed on top of the screen if participants were within the reward region (see figure), and of a red cross if they were not (not shown). The black solid line represents the hand-to-cursor discrepancy (the perturbation) for the SRT and FRT group across the task, and the grey dashed line represents the perturbation for the Gradual group only. Coloured lines represent group mean and shaded areas represent s.e.m. **(b)** Average reach angle of participants during baseline. **(c)** Average reach angle during the last 20 trials of the adaptation phase. The shaded grey area represents the region to be rewarded in the subsequent asymptote phase. **(d)** Average reach angle during the binary feedback (BF) block. The shaded grey area represents the rewarded region. **(e)** Success rate during the first 30 trials of the asymptote phase. **(f)** Success rate during the remainder of the asymptote phase (i.e. trial 31 to 200 of asymptote blocks). Each dot represents one participant. For the distribution plots, horizontal black lines are group medians and the shaded areas indicate distribution of individual values. SRT: short reaction time; FRT: fast reaction time. # p=0.059; *** p<0.001; ** p<0.01; * p<0.05.

During asymptotic performance, where participants were restricted to binary feedback, the SRT group showed a striking ability to maintain performance within the rewarded region whereas the two other groups clearly could not (figure 3d; H(2)=17.5, p<0.001, Bonferroni-corrected (see Methods), Tukey’s test p<0.001 vs FRT and p=0.001 vs Gradual). Next we compared success rates across groups for early BF trials (i.e. first 30 trials; figure 3e) and the remainder of BF trials (figure 3f) independently. Early success rates were significantly lower for the Gradual group compared to the SRT (H(2)=9.2, p=0.02, Bonferroni-corrected, Tukey’s test p=0.011), and a similar but nonsignificant trend was observed between the FRT and SRT groups (Tukey’s test p=0.059). The absence of a significant difference in early success rate between the FRT and SRT groups cannot be explained by average reach angles, as the FRT group actually express a larger decrease in reach angle during that timeframe compared to the Gradual group (figure 3a). Rather, the greater variability in reach angle within individuals in the FRT as opposed to the Gradual group is likely to cause this result (average individual variance; FRT: 47.5; Gradual: 18.9). However, success rate during the remaining trials reached significance for both the FRT and Gradual groups compared to the SRT group (H(2)=16.67, p<0.001, Bonferroni-corrected, Tukey’s test p<0.001 for both FRT and Gradual). Surprisingly, no dip in performance was observed for the SRT group in the early phase of the BF blocks, suggesting that informing participants of the perturbation and how to overcome it at the beginning of the experiment is sufficient to prevent this drop in reach angle.

Next, to ensure the low end adaptation reach angles expressed by the FRT group did not explain the low success rates, we removed every participant who expressed less than 15° reach angle at the end of the adaptation from each group (e.g. ^37^). Henceforth, we refer to those participants as non-adapters, as opposed to adapters. This procedure resulted in 1, 5 and 2 participants being removed in the SRT, FRT and Gradual groups, respectively. Performance for the adapters was fundamentally the same as the original groups (figure 4a), except for end adaptation reach angles, which were now all above 15° (figure 4b; SRT 17.0 ±1.2; FRT 16.9 ±1.2; Gradual 16.7 ±1.4). Specifically, the SRT-adapter group still showed a clear ability to remain in the rewarded region during binary feedback performance (asymptotic blocks), whereas the other two adapter groups could not (figure 4c; H(2)=14.0, p=0.002, Bonferroni-corrected, Tukey’s test p=0.028 vs FRT-adapter and p=0.001 vs Gradual-adapter). Because the full groups (i.e. non-Apdaters included) did not express a drop in success rate during early asymptote trials, we compared Adapters’ success rates during asymptote as a whole, rather than splitting them between early and late performance. The SRT-adapter group still displayed greater success than the Gradual-adapter group (figure 4d; H(2)=13.74, p=0.002, Bonferroni-corrected, Tukey’s test p<0.001). However, the difference between the SRT-adapter and the FRT-adapter group was now non-significant (Tukey’s test p=0.12). Despite this, the reach angle differences clearly show that successful binary performance remained strongly affected by one’s capacity to develop and express a strategy even for the successful adapters, as shown by the Gradual-adapter and FRT-adapter groups, respectively (figure 4a).

**Figure 4.**
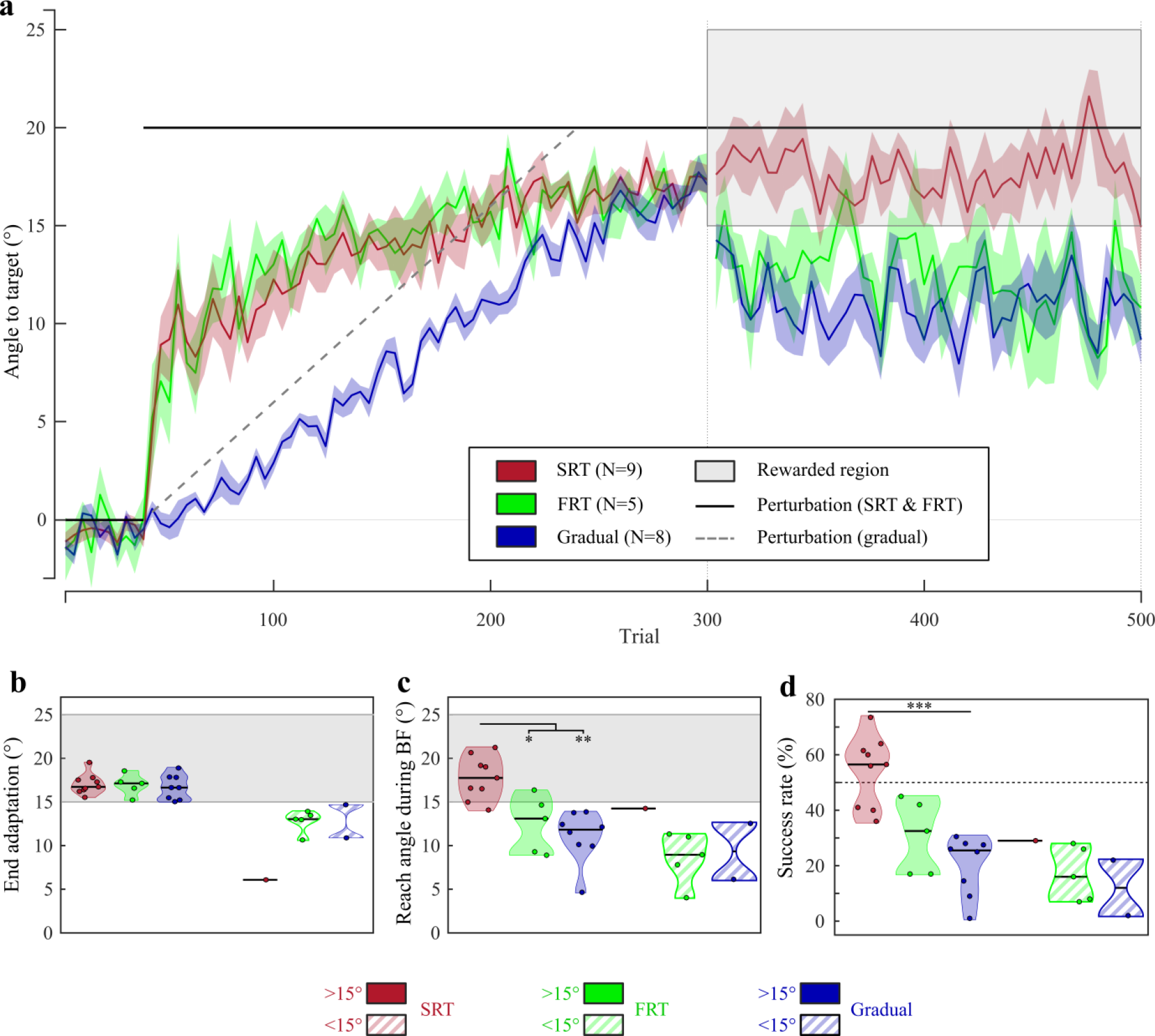
Performance of successful adapters during the forced RT task. **(a)** Reach angles with respect to target (°) of each group’s successful adapters exclusively. Values are averaged across epochs of 4 trials. Vertical bars represent block limits. The binary feedback consisted of a large green tick displayed on top of the screen if participants were within the reward region (see figure), and of a red cross if they were not (not shown). The black solid line represents the hand-to-cursor discrepancy (the perturbation) for the SRT and FRT group across the task, and the grey dashed line represents the perturbation for the Gradual group only. Coloured lines represent group mean and shaded areas represent s.e.m. **(b)** Average reach angle during the last 20 trials of the adaptation phase. The shaded area represents the region to be rewarded in the subsequent asymptote phase. **(c)** Average reach angle during the binary feedback (BF) block. **(d)** Success rate during the asymptote phase. The black dashed line represents 50% success rate. Each dot represents one participant. For the distribution plots, horizontal black lines are group medians and the shaded areas indicate distribution of individual values. >15° and <15° indicate the average reach angle during the end of the adaptation phase (i.e. adapter and non-adapter, respectively). SRT: short reaction time; FRT: fast reaction time. *** p<0.001; ** p<0.01; * p<0.05.

Finally, since trials were reinitialised if participants failed to initiate reaching movements within the allowed timeframe, we compared the average occurrence of these failed trials between the FRT and SRT groups (Supplementary figure S2c) to ensure any between-group difference cannot be explained by this. Both groups expressed similar amounts of failed attempts per trial (U=100, p=0.73). In addition, movement times were significantly faster across all blocks for the FRT group compared to the SRT group (Supplementary figure S2d; H(2)=11.78, p=0.005, Tukey’s test p=0.002), although they remained strictly under 400 ms for all groups as in the first experiment (figure 1c). RTs expressed by the Gradual group were between the SRT and FRT constraints (Supplementary figure S2a; Gradual group RT range 385 to 1610 ms).

Overall these findings demonstrate that preventing strategy use by restricting its expression or making participants unaware of the nature of the task results in the partial incapacity of participants to perform successfully during binary feedback performance. It should be noted, however, that performance does not reduce back to baseline entirely, as participants in both the FRT and Gradual groups are still able to express intermediate reach angle values of the order of 10 to 15°.

## Discussion

Previous work has led to the idea that BF induces recruitment of a model-free reinforcement system that strengthens and consolidates the acquired memory of a visuomotor displacement ^10,12,17^. Here, we investigated the role of explicit strategy-use in the context of BF, and our results suggest that it may have a more central role in explaining general BF-induced behaviours than previously expected. In the first experiment, the increased retention observed in the BF-Maintain group was suppressed if participants were told to “remove their strategy” (BF-Remove group). In the second experiment, preventing strategy-use by using a secondary task or preventing development of a strategy with a gradual introduction of the perturbation resulted in participants being unable to maintain accurate performance during BF blocks, suggesting that strategy-use is necessary for performing a BF reaching task.

The initial performance drop observed at the introduction of BF for both BF groups suggests that participants cannot immediately account for a visuomotor displacement they have already successfully adapted to ^12^. A possible explanation is that the cerebellar memory is not available anymore, most likely because removing VF results in a context change, which is known to prevent retrieval and expression of an otherwise available memory ^38–40^. Considering this, the restoration of performance observed after this dip could not be explained by recollection of the cerebellar memory, suggesting another mechanism took place. Two possible candidates to explain this drift back are model-free reinforcement ^10–12,17^ and strategy-use ^7,8,35^.

Reinforcement learning is usually considered to operate through experiencing success ^10,11^. It is thus difficult to argue for a reinforcement-based reversion to good performance during BF because participants in the trough of the dip do not experience a large amount of success, if any. Furthermore, participants experienced little “plateau” performance during the previous block, making formation of a model-free reinforcement memory unlikely, because it is considered a rather slow learning process as opposed to model-based reinforcement ^10,41^. On the other hand, both BF groups experienced a large amount of unexpected errors during this drop, which may promote a more strategy-based approach ^13–15,22^. In line with this, the SRT group in the forced RT task, which has been informed of the displacement and of the right strategy to counter it, does not express such dip when starting the BF block.

The forced RT task addresses this question more directly, and shows that impeding strategy-use with a secondary task ^30,32^ prevents participants from restoring performance over BF blocks, confirming our interpretation. Interestingly, both the FRT and Gradual groups do not show a return to baseline during asymptote. Likely, the FRT group is aware of the optimal strategy, and can partially express it, leading to these intermediate reach angles. Indeed, previous work on forced RT paradigms shows that adapting the constraints based on each individual’s baseline proficiency at this task more efficiently prevents strategy-use ^32^. On the other hand, the Gradual group was not informed of the optimal strategy, and thus would be expected to reach back to baseline‥ However, even in the presence of BF, the Gradual group shows a striking inability to find the optimal strategy, suggesting the lack of structural understanding of the task strongly impedes their exploration. This overall incapacity of the Gradual group to express an efficient explorative strategy is consistent with previous findings showing that rewarding success alone without providing any explanation of the task structure is not sufficient to make participants reliably learn an optimal strategy ^42^.

Previous studies employing the forced RT paradigm have shown it usually leads to slower learning rates during adaptation because participants can less easily apply a strategy from the beginning ^16,30,32^. In contrast, no such difference in learning rate was observed in our forced RT groups. This is possibly due to the difference in size of the perturbation between our study (20°) compared to others ^30,32^ (30°), making the explicit contribution potentially smaller during the adaptation phase ^7^.

Our findings qualitatively replicate results from a previous study employing a similar design ^12^. However, it should be noted that our paradigm differs in several ways. First, retention was assessed using feedback removal rather than visual error clamps, although there is evidence that both methods lead to quantitatively similar results ^43^. Second, our displacement was only 20° of amplitude and no additional displacement was introduced after the asymptote blocks. There is now a growing wealth of evidence that the cerebellum cannot account for more than 15 to 20° displacements ^32,36,44^, with the remaining discrepancy usually being accounted for through strategic re-aiming ^35^. Therefore, the absence of a second, larger displacement, if anything, should only result in a less strategy-based performance. Nevertheless, instructing participants to remove any strategy (Remove groups) resulted in a near-complete nullification of the binary feedback effect, suggesting it is mainly underlain by a simple re-aiming process. However, the Maintain instruction alone was not sufficient to produce this high retention profile, as the VF-Maintain group did not express it. We believe this can be explained in two ways. First, experiencing no feedback may result in a stronger context change for the VF groups compared to the BF groups, because the latter ones experienced the absence of VF during the asymptote blocks beforehand. Thus, this should lead to a stronger drop in reaching angle at the beginning of the no feedback trials for the VF groups, as observed here. Alternatively, the VF-Maintain group experienced 200 more trials with visual feedback at asymptote. Consequently, it is very likely that the cerebellar memory at the beginning of the no-feedback blocks was stronger ^11^, and the explicit contribution was less for this group compared to the BF-Maintain group ^7,16,35,45^. This would therefore result in the slow drop in reach angle observed during early no-feedback trials due to gradual decay of the cerebellar memory ^38,43,46^. Critically, both possibilities are not incompatible, and may well occur together.

A notable feature of retention performance is that both BF- and VF-Remove groups show a residual bias of around 5° in their reach angle in the direction of the displacement. Participants in the Remove conditions were not aware of this upon asking them after the experiment. This has been reliably observed in studies using no-feedback blocks to assess retention ^47,48^ (but see ^43^). Possible explanations include use-dependent plasticity-induced bias ^49,50^ or an implicit model-free reinforcement-based memory, although this study cannot provide any account toward one or the other. Note however that although the BF-Remove group expressed slightly more bias than its VF counterpart, this clearly did not reach statistical significance, meaning this cannot be explained by feedback type alone. Regardless, the implicit and lasting nature of this phenomenon makes it a promising focus for future research with clinical applications.

Overall, our findings all point toward a central role of strategy-use during BF-induced behaviours. In line with this, 14/54 participants had to be removed from the BF groups in the feedback-instruction task (experiment 1) because of poor performance in the asymptote blocks (see methods), suggesting that structural learning is required to perform accurately ^42^. This is again in line with the dip observed in the BF groups and the absence of dip in the (informed) SRT group. Our view is that implicit, model-free reinforcement takes a great amount of time and practice to form ^41,51^, and usually arises from initially model-based performance in behavioural literature ^18,52^, as illustrated by popular reinforcement models (e.g. DYNA ^53,54^). Two interesting possibilities are that 200 trials of BF alone are not sufficient to result in a strong, habit-like enhancement of retention ^52^, or that such behavioural consolidation must take place through sleep ^52,55^. Future work is required to address these hypotheses.

In conclusion, this study provides further insight into the use of reinforcement during motor learning, and suggests that successful reinforcement learning is tightly coupled to development and expression of an explicit strategy. Future studies investigating reinforcement during visuomotor adaptation should therefore proceed with care in order to map which behaviour is the consequence of actual implicitly reinforced memories or more explicit, strategic control.

## Methods

### Participants

80 participants (20 males) aged 18-37 (M=20.9 years) and 30 participants (11 males) aged 18-34 (M=22.1 years) were recruited for experiment one and two, respectively, and pseudo-randomly assigned to a group after providing written informed consent. All participants were enrolled at the University of Birmingham. They were remunerated either with course credits or money (£7.5/hour). They were free of psychological, cognitive, motor or auditory impairment and were right-handed. The study was approved by the local research ethics committee of the University of Birmingham and done in accordance to its guidelines.

### General procedure

Participants were seated before a horizontal mirror reflecting a screen above (refresh rate 60 Hz) that displayed the workspace and their hand position (figure 1a), represented by a green cursor (diameter 0.3 cm). Hand position was tracked by a sensor taped on the right hand index of each participant and connected to a Polhemus 3SPACE Fastrak tracking device (Colchester, Vermont U.S.A; sampling rate 120 Hz). Programs were run under MatLab (The Mathworks, Natwick, MA), with Psychophysics Toolbox 3 ^56^. Participants performed the reaching task on a flat surface under the mirror, with the reflection of the screen matching the surface plane. All movements were hidden from the participant’s sight. When each trial starts, participants entered a white starting box (1 cm width) on the centre of the workspace with the cursor, which triggered target appearance. Targets (diameter 0.5 cm) were 8 cm away from the starting position. Henceforth, the target position directly in front of the participant will be defined as the 0° position and other target positions will be expressed with this reference. Participants were instructed to perform a fast “swiping” movement through the target. Once they reached 8 cm away from the starting box, the cursor disappeared and a yellow dot (diameter 0.3 cm) indicated their end position. When returning to the starting box, a white circle displaying their radial distance appeared to help them get back into it.

### Task design

#### Experiment 1: feedback-instruction

For each trial, participants reached to a target located 45° counter-clock wise (CCW). Participants first performed a baseline block (60 trials) with veridical cursor feedback, followed by a 75 trials adaptation block in which a 20° CCW displacement was applied (figure 1b). In the following 2 blocks (100 trials each), participants either experienced the same perturbation with only BF, or with BF and VF. BF consisted of a pleasant sound selected based on each participant’s preference from a series of 26 sounds before the task, unbeknownst of the final purpose. When participants’ cursor reached less than 5° away from the centre of the target, the sound was played, indicating a hit; otherwise no sound was played, indicating a miss. For the BF group, no cursor feedback was provided, except for one “refresher” trial every 10 trials where VF was present. Participants in the VF group could see the cursor position at all times during the trial, along with the BF. Finally, participants went through 2 no-feedback blocks (100 trials each) with BF and VF completely removed. Before those blocks, participants were either told to “carry on” (“Maintain” group) or informed of the nature of the perturbation, and asked to stop using any strategy to account for it (“Remove” group). Therefore, we had four groups in a 2x2 factorial design (BF versus VF and Maintain versus Remove). Finally, if a trial’s reaching movement duration was greater than 400 ms or less than 100 ms long, the starting box turned red or green, respectively, to ensure participants performed ballistic movements, and didn’t make anticipatory movements. Participants who expressed a success rate inferior to 40% during asymptote blocks were excluded (BF-Remove N=6; BF-Maintain N=8). Although this exclusion rate was high, it was crucial to exclude participants who were unable to maintain asymptote performance in order to reliably measure retention.

#### Experiment 2: forced RT

In this experiment, participants were forced to perform the same reaching task at slow (SRT) or fast reaction times (FRT), the latter condition preventing strategy-use by enforcing movement initiation before any mental rotation can be applied to the motor command ^30,33^. A third group (Gradual) also performed the task with no RT constraints.

In the SRT/FRT groups, for each trial, entering the starting box with the cursor triggered a series of five 100 ms long pure tones (1 kHz) every 500 ms (figure 1c). Before the fifth sound, a target appeared at one of four possible locations equally dispatched across a span of 360° (0-90-180-270°). Participants were instructed to initiate their movement exactly on the fifth tone (figure 1c). Targets appeared 1000 ms (SRT) or 200 ms (FRT) before the beginning of the fifth tone. Movement initiations shorter than 130 ms are likely anticipatory movements ^31^, and explicit strategies start to be difficult to express under 300 ms ^30,32^. Therefore, in both conditions, movements were successful if participants exited the starting box between 70 ms before the start of the fifth tone and the end of the fifth tone, that is, from 130 ms to 300 ms after target appearance in the FRT condition. If movements were initiated too early or too late, a message "too fast" or "too slow" was displayed and the cursor did not appear upon exiting the starting box. The trial was then reinitialised and a new target selected. Finally, if participants repeatedly missed movement initiation, making trial duration over 25 seconds, RT constraints were removed, to allow trial completion before cerebellar memory time-dependent decay ^43,46,57^. Participants in the SRT and FRT groups were informed of the displacement and of the optimal strategy to counter it, to ensure that any effect was related to expression, rather than development of a strategy. They were also instructed to attempt using the optimal strategy as much as possible when sensible, but not at the expense of the secondary RT task, so as to preserve the pace of the experiment and prevent time-dependent memory decay.

To attain proficiency in the RT task, SRT and FRT participants performed a training block (pseudo-random order of VF and BF trials) of at least 96 trials, or until they could initiate movements on the fifth tone reliably (at the first attempt) at least for 75% of the previous 8 trials. All participants achieved this in 96 to 157 trials. Once this was achieved, participants first performed a 40 trials baseline (figure 1d), followed by introduction of a 20° CCW displacement for 260 trials. Participants then underwent a 200-trials long asymptote block with only BF (1 “refresher” trial every 10 trials). The BF consisted of a green tick or a red cross if participants hit or missed the target, respectively. Visual BF was used to prevent interference with the tones presented to manipulate RTs. The Gradual group underwent the same schedule, except that no tone or RT constraint were used, and the perturbation was introduced gradually from the 41^st^ to the 240^th^ trial of the first block (increment of 0.4°/trial) occurring independently for each target. This ensured participants experienced as few large errors as possible to prevent awareness of the perturbation and therefore strategy-use. After the experiment, participants in the Gradual group were informed of the displacement, and subsequently asked if they noticed it. If they answered positively, they were asked to estimate the size of the displacement.

### Data analysis

All data and analysis code is available on our open science framework page (osf.io/hrgzq). All analyses were performed in MatLab. We used Lilliefors test to assess whether data were parametric, and we compared groups using Kruskal-Wallis or Wilcoxon signed-rank tests when appropriate, as most data were non-parametric. Post-hoc tests were done using Tukey’s procedure. As we analysed the data from experiment two twice (figure 3 and 4), success rates and reach angles during asymptote were Bonferroni-corrected with corrected p-values (multiplied by 2).

Learning rates were obtained by fitting an exponential function to adaptation block reach angle curves with a non-linear least-square method and maximum 1000 iterations (average R^2^ = 0.86 ±0.14 for feedback-instruction task and R^2^ = 0.58 ±0.26 for forced-RT task):

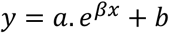

where *y* is the hand direction for trial *x*, *a* is a scaling factor, *b* is the starting value and *β* is the learning rate. Reach angles were defined as angular error to target of the real hand position at the end of a movement. Trials were considered outliers and removed if movement duration was over 400 ms or less than 100 ms, end point reach angle was over 40° off target, and for the SRT and FRT groups in the forced-RT task, if failed initiation attempts continued for more than 25 sec. In total, outliers accounted for 3755 trials (8%) in the feedback-instruction task and 1013 trials (6%) in the forced-RT task.

Even though 4 targets were used during the forced-RT task, trials were reset and a new random target was selected every time participants failed to initiate movements on the 5^th^ tone. Therefore, all possible target positions would not be represented for each epoch and analysis was done without using epochs.

## Acknowledgements

We thank Raphael Schween for helpful discussions on interpretation of the data. This work was supported by the European Research Council grant MotMotLearn 637488.

## Additional information

The authors declare no competing financial interests.

## Author contributions

O.C., P.J.H. and J.M.G. designed the experiments, O.C. implemented and ran the experiments, O.C. and P.J.H. analysed the data, O.C., P.J.H. and J.M.G. interpreted the results, O.C. wrote the paper, O.C., P.J.H. and J.M.G. approved the final version of the manuscript.

## Supplementary figures

**Supplementary figure S1.**
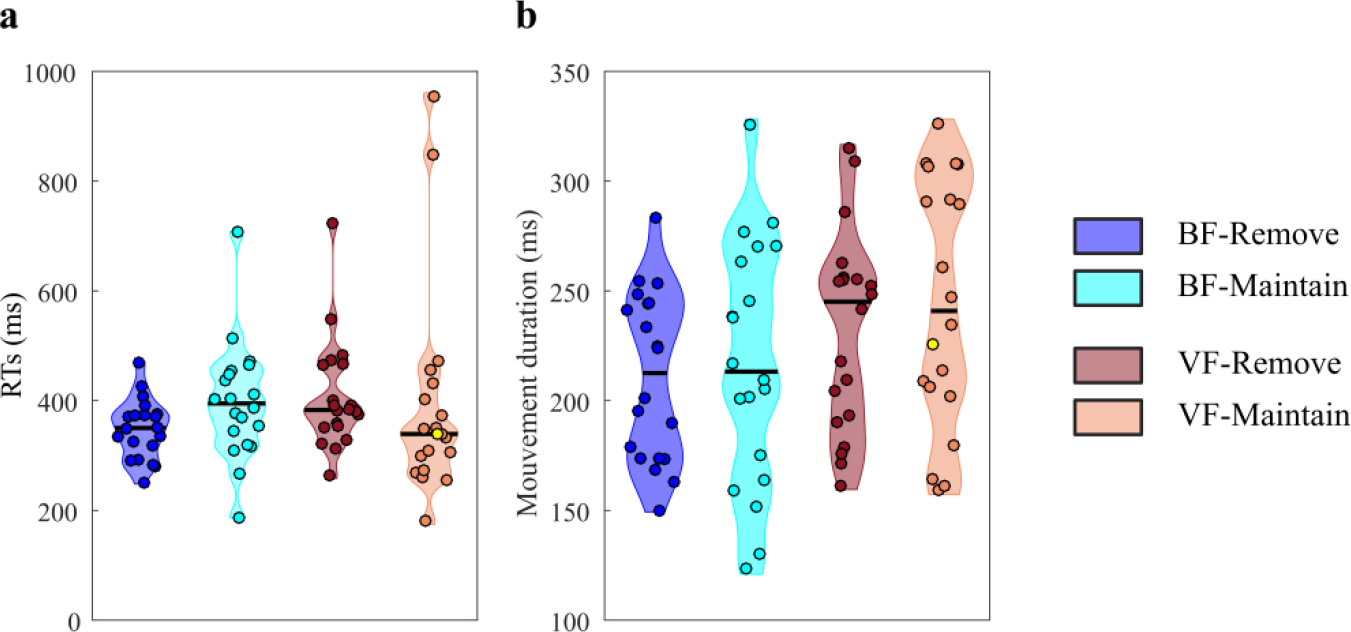
Experiment 1: feedback-instruction. **(a)** Average reaction times of participants during the asymptote phase. **(b)** Average movement duration of participants during the asymptote phase. Each dot represents one participant. The yellow dot represents the same participant across all plots (the same participant as figure 2). Black lines are group medians and the shaded areas indicate distribution of individual values.

**Supplementary figure S2.**
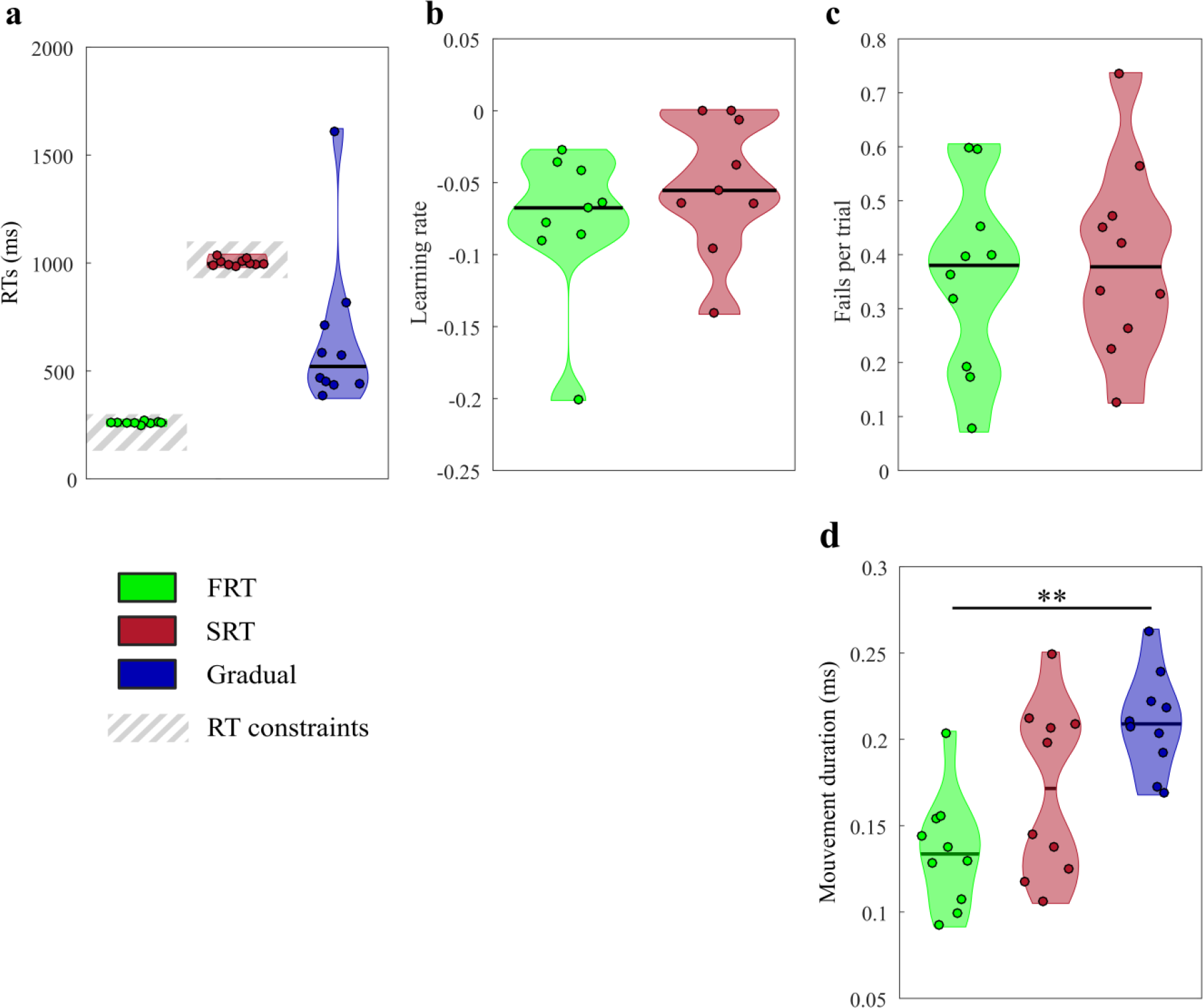
Experiment 2: forced RT. **(a)** Average reaction times of participants throughout the task. **(b)** Average number of failures per trial to initiate movements within the constrained timeframe throughout the task. **(c)** Average movement duration of participants throughout the task. **(d)** Learning rates during the adaptation phase. Each dot represents one participant. Black lines are group medians and the shaded areas indicate distribution of individual values. SRT: short reaction time; FRT: fast reaction time. ** p<0.01.

